# T cell epitope mapping reveals immunodominance of evolutionarily conserved regions within SARS-CoV-2 proteome

**DOI:** 10.1101/2024.10.23.619918

**Authors:** Cansu Cimen Bozkus, Matthew Brown, Leandra Velazquez, Marcus Thomas, Eric A. Wilson, Timothy O’Donnell, Denis Ruchnewitz, Douglas Geertz, Yonina Bykov, Julia Kodysh, Kasopefoluwa Y. Oguntuyo, Vladimir Roudko, David Hoyos, Komal D. Srivastava, Giulio Kleiner, Hala Alshammary, Neha Karekar, Christopher McClain, Ramya Gopal, Kai Nie, Diane Del Valle, Daniela Delbeau-Zagelbaum, Denise Rodriguez, Jessica Setal, The Mount Sinai COVID-19 Biobank Team, Emily Carroll, Margrit Wiesendanger, Percio S. Gulko, Alexander Charney, Miriam Merad, Seunghee Kim-Schulze, Benhur Lee, Ania Wajnberg, Viviana Simon, Benjamin D Greenbaum, Diego Chowell, Nicolas Vabret, Marta Luksza, Nina Bhardwaj

## Abstract

As SARS-CoV-2 variants continue to emerge capable of evading neutralizing antibodies, it has become increasingly important to fully understand the breadth and functional profile of T cell responses to determine their impact on the immune surveillance of variant strains. Here, sampling healthy individuals, we profiled the kinetics and polyfunctionality of T cell immunity elicited by mRNA vaccination. Modeling of anti-spike T cell responses against ancestral and variant strains of SARS-CoV-2 suggested that epitope immunodominance and cross-reactivity are major predictive determinants of T cell immunity. To identify immunodominant epitopes across the viral proteome, we generated a comprehensive map of CD4^+^ and CD8^+^ T cell epitopes within non-spike proteins that induced polyfunctional T cell responses in convalescent patients. We found that immunodominant epitopes mainly resided within regions that were minimally disrupted by mutations in emerging variants. Conservation analysis across historical human coronaviruses combined with *in silico* alanine scanning mutagenesis of non-spike proteins underscored the functional importance of mutationally-constrained immunodominant regions. Collectively, these findings identify immunodominant T cell epitopes across the mutationally-constrained SARS-CoV-2 proteome, potentially providing immune surveillance against emerging variants, and inform the design of next-generation vaccines targeting antigens throughout SARS-CoV-2 proteome for broader and more durable protection.

**One Sentence Summary:** Polyfunctional CD8+ and CD4+ T cells directed against SARS-CoV-2 target mutationally constrained regions of the viral proteome.

## INTRODUCTION

Since its initial emergence in December 2019, the severe acute respiratory syndrome coronavirus 2 (SARS-CoV-2) genome has consistently undergone mutations, resulting in new variants. Some of these variants, Alpha, Beta, Gamma, Delta and Omicron, spread rapidly, causing outbreaks, and are designated as “variants of concern (VOC)”. VOC are marked by their enhanced transmissibility, effectively outcompeting their predecessors due to increased viral fitness and escape from immune recognition (*1*). Although the COVID-19 vaccines are continuously adapted to target prevalent VOC, by the time they are commonly accessible, the next variant will likely have emerged, compromising the vaccine’s efficacy. This race between the new variants and effective vaccine development warrants the design of next-generation vaccines that can provide unfettered protection against future SARS-CoV-2 variants.

Coordinated antigen-specific B and T cell responses, induced by vaccination and/or infection, are imperative to clearing RNA viruses (*2*, *3*). Accordingly, the role of neutralizing antibodies in preventing SARS-CoV-2 infection (*4*) and improving disease outcomes has been well documented (*5*). However, antibody titers wane over time (*6*) and VOC frequently evade neutralization by antibodies (*7–9*). Despite reports of certain mutations evading T cell recognition (*10*, *11*), there is accumulating evidence that demonstrates preservation of overall T cell responses against VOC by targeting conserved prevalent epitopes (*12*) or cross-recognizing mutated epitopes (*13*). Notably, T cell responses are critical for controlling SARS-CoV-2 infection and SARS-CoV-2-specific CD4+ and CD8+ T cells correlate with durable immune protection and reduced disease severity (*14–16*). Corroborating the durability of T cell immunity, COVID-19 vaccines have been shown to induce stem-like memory T cells persisting more than 6 months after vaccination (*17*). Additionally, in preclinical studies using transgenic mice infected with SARS-CoV-2 (*18*) and rhesus macaques administered intranasal SARS-CoV-2 vaccines, CD8+ T cells alone were shown to be adequate for viral clearance even in the absence of humoral immunity (*19*). Furthermore, in patients receiving B cell-depleting therapies, vaccines elicited sustained T cells responses against SARS-CoV-2, providing protection from severe disease (*20*, *21*). Therefore, T cells may be key to mediating long-lasting, protective immunity against SARS-CoV-2 amid emerging variants that can evade antibody responses.

Detailed interrogation of T cell specificities from patient cohorts exposed to SARS-CoV-2 has demonstrated the induction of T cell responses against a range of viral proteins, among which spike (S), nucleocapsid (N), membrane (M), ORF1, and ORF3 appear to be dominant targets (*22–25*). Additionally, studies of unexposed individuals identified preexisting memory T cells recognizing SARS-CoV-2 sequences (*22*, *26*, *27*), which are reported to be cross-reactive T cells derived from previous exposure to common cold viruses (HCoVs) (*28*). HCoVs share partial sequence homology to SARS-CoV-2, across both structural and non-structural proteins, (*29*) and circulate widely, having infected a significant proportion of the population (*30*, *31*). A potential protective role for HCoV-derived cross-reactive T cells has also been suggested in individuals subsequently exposed to SARS-CoV-2 (*32*). Thus, incorporating immunogenic T cell epitopes from conserved regions of the SARS-CoV-2 proteome into immunization strategies may offer durable and comprehensive protection, as reinforcement against emerging variants that escape humoral immunity (*33*).

Here, we first investigated the specificities of T cell responses towards immunogenic regions in S that are subject to mutational events in SARS-CoV-2 VOC in cohorts of vaccinated individuals. Our kinetics studies confirmed that T cells were induced early on after mRNA vaccination with a high frequency of polyfunctional populations able to contribute to coordinated adaptive immunity. To understand the determinants governing the preservation of T cell responses and viral immune escape, we extensively mapped the functional responses to mutated S epitopes revealing that polyclonal T cell populations were preserved and cross-reactive against variant antigens. Additionally, because internal antigens also serve as potent targets of T cell immunity and experience less antibody-driven mutational pressure compared to S, we conducted one of the largest experimental efforts to-date for combined CD4+ and CD8+ T cell epitope mapping of non-S proteins. Our comprehensive approach revealed highly immunogenic, dominant T cell epitopes across a cohort of convalescent patients, eliciting conserved cellular immune responses, which could serve as candidate epitopes for incorporation into vaccine designs effective against emerging SARS-CoV-2 variants and potentially pan-coronavirus vaccine strategies.

## RESULTS

### Adaptive immune responses elicited by SARS-CoV-2 mRNA vaccination

To study SARS-CoV-2 mRNA vaccine-induced adaptive immunity, we collected longitudinal blood samples from 15 healthy donors, who received either mRNA-1273 (by Moderna) or BNT162b2 (by Pfizer/BioNTech) vaccine series (table S1). Samples were collected before vaccination (V0), 14 days after 1st dose (V1D14), 7 days after 2nd dose (V2D7), and 14 days after 2nd dose (V2D14) (Fig. 1A). We assessed the kinetics of the neutralizing antibody responses, a major correlate of protection from SARS-CoV-2 (*4*), using a pseudotyped vesicular stomatitis virus (VSVΔG-Rluc) expressing the D614G SARS-CoV-2 spike (S) protein.

**Fig. 1.**
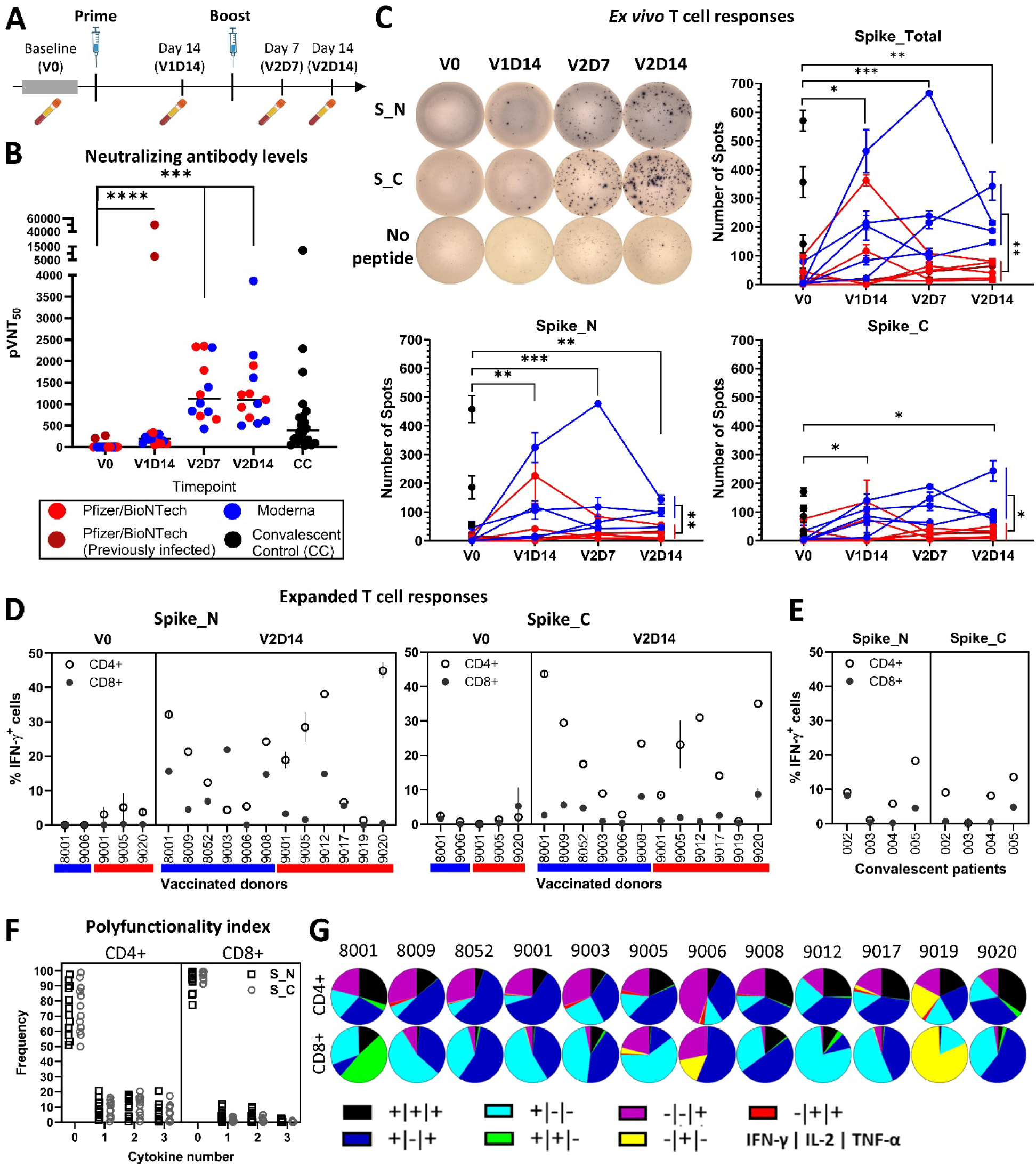
SARS-CoV-2 mRNA vaccine-induced adaptive immunity. **(A)** Peripheral blood samples were collected from individuals receiving COVID-19 mRNA vaccines, mRNA-1273 (by Moderna) or BNT162b2 (by Pfizer/BioNTech) longitudinally: before vaccination (V0), 14 days after 1st dose (V1D14), 7 days after 2nd dose (V2D7) and 14 days after 2nd dose (V2D14), and from convalescent COVID-19 patients, who were not vaccinated. **(B)** Serum neutralization capacity was assessed by a pseudotype particle (pp) infection system, VSVΔG-Rluc bearing the SARS-CoV-2 D614G spike glycoprotein targeting 293T cells stably expressing ACE2 and TMPRSS2. 4-point nonlinear regression curves were used to calculate 50% pseudovirus neutralization titers (pVNT50) for vaccine recipients (n=12) and convalescent patients (n=20). Horizontal lines denote median pVNT50 values. Peripheral blood mononuclear cells (PBMCs, 2×10^5^ cells/well) were stimulated with pools of overlapping peptides, 15mers with 5 amino acid offsets, spanning the N-terminal (Spike_N or S_N) or C-terminal half (Spike_C or S_C) of Spike protein for 24 h and IFN-γ secretion was measured by ELISPOT. **(C)** Representative ELISPOT wells from a vaccinated donor (top left) and summary of ELISPOT data (n=11 vaccinated donors, BNT162b2 recipients in shown red, mRNA-1273 recipients in blue, and n=5 convalescent patients, shown in black), where Spike_Total (top right) is the sum of spots acquired by Spike_N (bottom left) and Spike_C (bottom right) peptide pools. T cells were expanded following stimulation with Spike_N and Spike_C peptide pools. Antigen-specific cytokine production by expanded T cell subsets, CD4+ or CD8+, were measured by intracellular staining by flow cytometry. IFN-γ production by Spike-specific T cells **(D)** in vaccinated donors and **(E)** convalescent patients. **(F)** Polyfunctionality of Spike-specific T cells at V2D14 as demonstrated by % of T cells (y axis) co-expressing effector cytokines: IFN-γ, TNF-α, and IL-2 (x axis). **(G)** Distribution of Spike-specific (Spike_Total) T cell responses in each vaccinated donor at V2D14. Spot numbers and cytokine+ cell frequencies were demonstrated after background subtraction. Statistical significance (p < 0.05) was evaluated by Wilcoxon matched-pairs test by comparing vaccination timepoints and Welch’s t-test was used for comparing T cell responses elicited by mRNA-1273 vs BNT162b2.

Neutralizing antibody titers were significantly increased after a single dose of mRNA vaccine and continued to increase after the second dose exceeding the titer levels detected in most unvaccinated convalescent control (CC) patients (Fig. 1B and fig. S1). Notably, 2 of the individuals in the vaccination cohort were previously infected with SARS-CoV-2. A single dose of mRNA vaccine was sufficient to robustly boost neutralizing antibody titers in these individuals (Fig. 1B).

Next, we interrogated the kinetics of vaccine-induced T cell responses, another key correlate of protection from SARS-CoV-2 (*14*). Peripheral blood mononuclear cells (PBMCs) collected from vaccinated donors at V0, V1D14, V2D7, and V2D14 were stimulated in an *ex vivo* IFN-γ ELISpot assay with pooled 15mer overlapping peptides (OLPs) spanning the N- and C-terminal halves of Wuhan-1 S protein, Spike_N and Spike_C, respectively (table S2). mRNA vaccination significantly induced anti-S T cell responses at all timepoints tested, demonstrating *in vivo* priming. However, there was no significant change in the magnitude of T cell responses across post-vaccine timepoints tested (Fig. 1C). Notably, the magnitude of anti-S T cell responses elicited by mRNA-1273 was significantly greater than those elicited by BNT162b2 vaccine (Fig. 1C). This may be due to different antigen loads and dosing schedules of the two mRNA vaccines (*34*). Similar differences between mRNA-1273 and BNT162b2-induced humoral responses were previously reported (*34*).

To study the abundance and functionality of vaccine-induced T cells more robustly, we expanded Wuhan-1 S-specific T cells in pre- and post-vaccination blood collected at V0 and V2D14, respectively. T cells were stimulated with Spike_N and Spike_C OLP pools and antigen-specific T cells were expanded for 10 days before measuring effector cytokine production by intracellular flow cytometry (*35*). Vaccination induced robust anti-S T cell responses in all donors tested.

Although less frequently, anti-S T cells were detected at V0 in some donors (Fig. 1D). This can be due to *in vitro* priming of naïve T cells (*35*) and/or expansion of cross-reactive T cells. mRNA vaccines could elicit both CD4+ and CD8+ anti-S T cells; however, reactive T cells were predominantly CD4+ in most of the donors (Fig. 1D). This was consistent with our observations in unvaccinated convalescent donors (CPC cohort, table S1), where infection-induced anti-S T cells were also predominantly CD4+ (Fig. 1E). As expected by their unique major histocompatibility complex (MHC) molecules (table S1) and T cell repertoires, the abundance and CD4:CD8 subset distributions of anti-S T cells varied across donors. Importantly, vaccines could induce polyfunctional CD4+ and CD8+ anti-S T cells secreting IFN-γ, TNF-α and IL-2 (Fig. 1F and G), further supporting the role of mRNA vaccination in establishing an effective T cell immunity against SARS-CoV-2. Induction of polyfunctional anti-S T cells was similar between mRNA-1273 and BNT162b2 recipients (fig. S2).

### Cross-reactivity of vaccine-induced adaptive immunity against SARS-CoV-2 variants

Breakthrough infections caused by emerging variants of concern indicate an impaired ability of the vaccine-induced immunity to recognize SARS-CoV-2 variants. To assess the kinetics of Wuhan-1 S-directed antibody binding to variant strains, we utilized sera collected at V0, V1D14, V2D7, and V2D14 in a Luminex binding assay with multiple spike receptor binding domain (RBD) constructs. Antibody recognition of ancestral and Alpha (B.1.1.7) strain RBD was comparable across all timepoints tested, while there was a reduction in the recognition of Beta (B.1.351) and Gamma (P.1) RBD (Fig. 2A), consistent with other studies showing limited antigenic change with Alpha (*36*) and reduced viral neutralization with Beta and Gamma variants (*37*). The reduced recognition of Beta and Gamma RBD was even more pronounced in convalescent patients and was at least in part mediated by E484K mutation found in both strains with potential contributions by the K417N mutation found in Beta, Delta and Omicron strains (Fig. 2A).

**Fig. 2.**
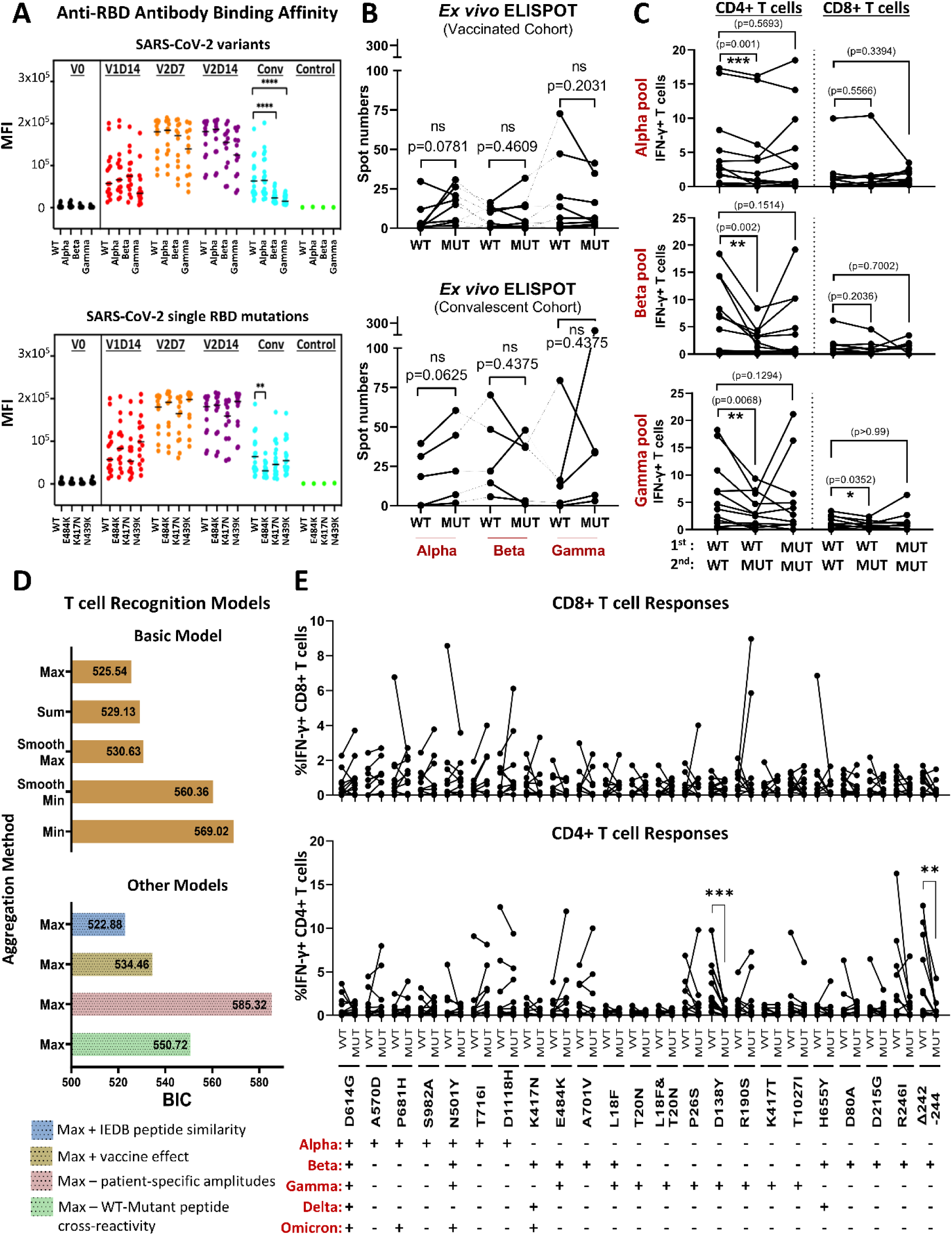
Adaptive immune recognition of SARS-CoV-2 variants. **(A)** Antibody reactivity from vaccinated (before vaccination (V0), 14 days after 1st dose (V1D14), 7 days after 2nd dose (V2D7) and 14 days after 2nd dose (V2D14)), convalescent (conv), or pre-COVID-19 (control) serum to RBD were assessed by Luminex antibody binding assay where MagPlex-C Microspheres Regions were conjugated to recombinant wild-type (WT, Wuhan-1) and mutant RBD constructs (Alpha [N501Y], Beta [N501Y/K417N/E484K], and Gamma [N501Y/K417T/E484K] with mean fluorescence intensity (MFI) used as a readout for binding affinity. **(B)** Summary of ELISPOT data: peripheral blood mononuclear cells (PBMCs, 2×10^5^ cells/well) from vaccinated donors (V2D14) or from convalescent patients were stimulated with pooled peptides covering the mutations found in Alpha, Beta and Gamma variants (listed in (E)) or the corresponding WT sequences for 24 h and IFN-γ secretion was measured by ELISPOT. **(C)** V2D14 T cells from vaccinated donors were stimulated with variant or WT peptide pools and expanded prior to being re-stimulated with either the initial stimulation peptide pool (WT→WT, Mut→Mut) or the variant pool to measure cross-reactivity (WT→Mut). Antigen-specific cytokine production by expanded T cell subsets, CD4+ or CD8+, was measured by intracellular staining by flow cytometry. **(D)** Bars show Bayesian Information Criterion (BIC) values for different models of T cell reactivity shown in C, where lower BIC is better. The basic model includes the effect of peptide pool stimulations and WT-Mutant peptide cross-reactivity. Different peptide dominance models are shown on the y axis, which correspond to the 9mer aggregation function used. Performance of other models was also measured, blue: accounting for sequence similarity with IEDB epitopes, brown: including vaccination effect as the initial stimulation event, pink: excluding the effect of patient-specific amplitudes, green: excluding the effect of cross-reactivity between peptides from 1^st^ and 2^nd^ stimulation events. **(E)** Deconvolution of IFN-γ production by T cells in response to individual mutations within the peptide pools tested in C (WT→WT vs Mut→Mut). Statistical significance (p < 0.05) was evaluated by Wilcoxon matched-pairs test.

Cross-reactive T cell responses are also important for controlling infections with SARS-CoV-2 variant strains, especially given the diminished neutralizing antibody control. To study cross-reactive T cell responses against S mutations identified in variants of concern, we designed 15mer OLPs spanning a maximum of 14 upstream and downstream amino acid (aa) sequences surrounding each mutated aa. Then we pooled the WT (Wuhan-1) and mutant OLPs corresponding to each variant tested, namely Alpha, Beta, and Gamma (table S2 and fig. S3). Using these pools, we stimulated PBMCs collected from vaccinated (V2D14) and convalescent (unvaccinated) individuals in an *ex vivo* IFN-γ ELISpot assay to measure the T cell recognition of WT and variant S. Both vaccine and infection-primed Wuhan-1 S-specific T cells recognized mutated S in Alpha, Beta and Gamma variants at a similar capacity *ex vivo* (Fig. 2B). These observations are in accordance with previous studies demonstrating preservation of overall T cell responses against early variants in *ex vivo* assays (*12*, *13*). However, multiple studies have reported that certain S mutations may lead to escape from T cell recognition (*10*, *11*). It is possible that *ex vivo* analysis of T cell cross-reactivity may not capture nuanced differences in the antigenicity of S mutations since total anti-S T cells comprise a small proportion among PBMCs and the mutated S epitopes tested here are only a subset of many immunogenic S epitopes engendering a wide breadth of T cell responses (*38*). Therefore, we aimed to study the antigenicity and cross-recognition of S mutations in clonally enriched T cell populations specific to these epitope regions. We expanded T cells from vaccinated donors collected at V2D14 stimulated with variant peptide pools and the corresponding WT pools, as described for *ex vivo* analysis. To measure antigen-specific effector responses, expanded T cells were re-stimulated with either the WT or mutant peptides with which they were initially stimulated. In addition, expanded WT T cells were re-stimulated with the corresponding mutant epitopes to directly measure the cross-recognition of mutant S epitopes by Wuhan-1 S-specific T cells. In accordance with total S-specific T cell immunity measured in fig. 1D, WT and mutated S-specific reactive T cells were predominantly CD4+ (Fig. 2C). Both WT and mutated S in Alpha, Beta, and Gamma variants induced overall T cell reactivity similarly (Fig. 2C). However, the direct measurement of cross-recognition revealed a significantly diminished reactivity of WT T cells against variant S (Fig. 2C).

To evaluate the underlying parameters of T cell immunity, we modeled anti-S T cell responses against ancestral and variant strains of SARS-CoV-2 measured in fig. 2C. Our basic model, detailed in the methods section, considers T cell recognition as a two-step process: 1) peptide binding and presentation by MHC (pMHC) and 2) binding of pMHCs by specific T cell receptors (TCRs), and includes the effects of initial stimulation, re-stimulation, and peptide cross-reactivity, as well as patient specific amplitude of immune response. Our pMHC calculations utilize prediction algorithms, which perform better for MHC-I than MHC-II binding (*39*). Therefore, we focused on CD8+ T cell responses. We computed a recognition score for each 9mer within the peptide pools over the MHC-I set for each donor. To evaluate the contribution of 9mers to overall pool level response, we applied different 9mer aggregating functions, including “*max*”, “*sum*”, and biologically unrealistic “*min*” as a control. The *max* function assumes that the most immunogenic 9mer in a pool is largely responsible for the observed T cell response, while the *sum* function assumes all 9mers contribute to the pool-level response in proportion to their relative immunogenicity. To compare different aggregating functions, we derived the Bayesian information criterion (BIC) values for each, where a lower value is preferred. The *max* function showed the best performance (Fig. 2D and fig. S4), suggesting that anti-S T cell responses are primarily mounted by immunodominant epitopes. Expectedly, the control *min* function, which assumes the least immunogenic epitope would drive the observed response, performed poorly. Further supporting the immunodominance model over the additive model, the *max* and *sum* functions performed similarly since the recognition scores of 9mers in a peptide pool were not uniform but typically dominated by a small number of 9mers with negligible scores for the remaining 9mers (Fig. 2D and fig. S5).

For the immunodominance model, we also compared extended and partial recognition models. Introducing a term accounting for the sequence similarity of our test epitopes and immunogenic epitopes in the immune epitope database (IEDB) slightly improved model performance in accordance with previous reports attributing this approach to increased TCR response (*40*, *41*). Removing patient-specific immune response amplitudes by setting them all to the same optimized value displayed significantly reduced performance (Fig. 2D), highlighting the heterogeneity of anti-S T cell responses observed across different donors. Our basic model accounts for cross-reactivity of peptides, computing their sequence-based distance and resulting impact on binding strength to the same TCR (*42*). Because all patients in our cohort were vaccinated, we included the vaccine as a zeroth stimulation event, introducing additional cross-reactivity terms with the complete Wuhan-1 S 9mers. However, we did not observe any improvement in the model’s performance, likely because the experimental data reflects only a small subset of anti-S T cells and *in vitro* expansion impacts clonal dynamics. To evaluate the role of cross-reactivity in variant S recognition by WT T cells more directly, we excluded the effect of initial stimulation and consequently observed a decreased performance (Fig. 2D).

Together these results suggest that immunodominant epitopes are key inducers of anti-S T cell immunity and that the overall anti-S T cell responses are preserved against mutant variants. As the direct measurement of cross-recognition suggested a diminished reactivity of WT CD4+ T cells against variant S, we investigated whether any specific S mutation may lead to escape from T cell recognition. Stimulation of vaccinated donor (V2D14) T cells with peptides spanning individual mutations found in variant S and the corresponding WT demonstrated a significant decrease across the population in the recognition of mutations P26S and R246I by CD4+ T cells (Fig. 2E). Although the impact of individual variant spike mutations on the T cell immunity is likely compensated by the wide breadth of T cell responses elicited in vaccinated individuals, the reduced cross-recognition of variant S and immune escape at certain mutant S epitopes prompt the inclusion of non-spike epitopes in future vaccine designs to further boost T cell immunity against SARS-CoV-2.

### T cell epitope mapping of non-spike SARS-CoV-2 proteins in convalescent patients

To evaluate the immunogenicity of non-spike epitopes across SARS-CoV-2 proteome, we synthesized overlapping 15mer peptides covering the entire nucleocapsid (N) and selected regions in other proteins (Fig. 3A and table S2). Selected regions were determined by prioritizing those enriched in interactions between epitopes predicted to be strong binders (*43*) and MHC alleles frequent across different races (*44*) (fig. S6). We measured T cell responses in unvaccinated convalescent donors (ATLAS cohort, table S1) utilizing *ex vivo* IFN-γ ELISpot and T cell expansion assays (Fig. 3B). Both S and non-S SARS-CoV-2 proteins elicited robust T cell responses *ex vivo* in convalescent patients (Fig. 3C) compared to healthy donor controls sampled prior to COVID-19 pandemic (fig. S7). The distribution of memory T cell responses against each protein varied across patients with S, N, membrane (M) and ORF7-directed responses being the most potent (Fig. 3D). Expansion assays confirmed the immunogenicity of non-S proteins and demonstrated that like anti-S T cells, non-S-specific T cells were also predominantly CD4+ (Fig. 3, E and F).

**Fig. 3.**
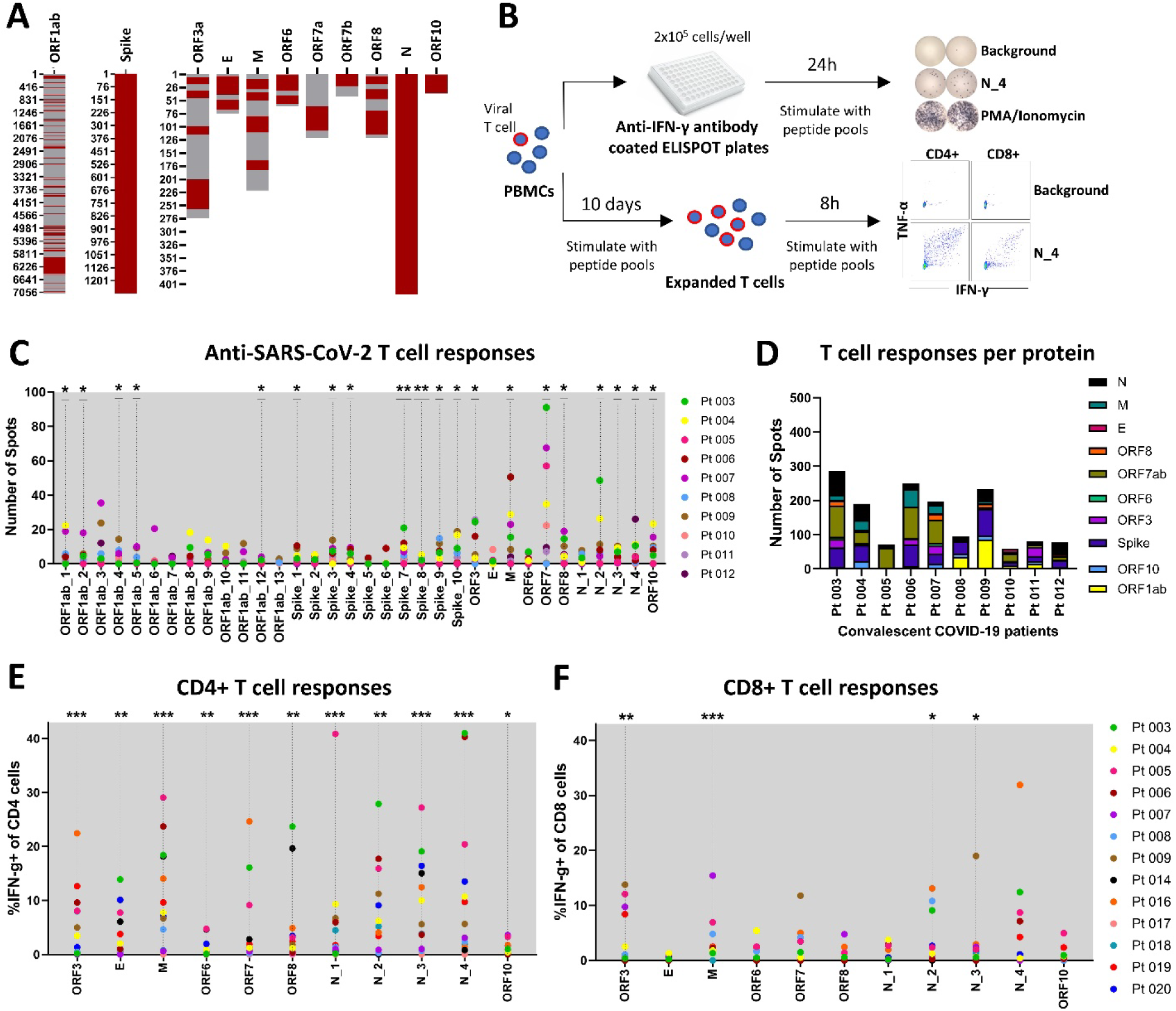
T cell responses against non-spike SARS-CoV-2 proteins. **(A)** Schema demonstrating peptide selection for each protein. The y axis denotes amino acid residue number. Tested regions are shown in red. **(B)** Experimental design summary. Peripheral blood mononuclear cells (PBMCs, 2×10^5^ cells/well) from convalescent, unvaccinated patients (n=10) were stimulated for 24 h with pools of overlapping peptides, 15mers with 5 amino acid offsets, spanning each protein as indicated on the x axis. Pools contained no more than 25 peptides. IFN-γ secretion was measured by ELISPOT. Summary of ELISPOT data showing total T cell responses **(C)** per peptide pool and **(D)** per patient. T cells from the same convalescent, unvaccinated patient cohort (n=15) were expanded following stimulation with non-spike peptide pools. Antigen-specific cytokine production (IFN-γ+) by expanded T cell subsets, **(E)** CD4+ or **(F)** CD8+, was measured by intracellular staining by flow cytometry. Each dot corresponds to a patient. Normalized values were shown. Statistical significance (p < 0.05) was evaluated by the Wilcoxon matched-pairs test comparing DMSO vs peptide stimulation.

Since our modeling of anti-S T cell responses suggested that T cell immunity was primarily elicited by immunodominant epitopes, we evaluated the presence of immunodominant regions within non-S proteins. We stimulated convalescent donor T cells with individual 15mer peptides constituting the peptide pools used in fig. 3. Epitope mapping revealed immunogenic regions across non-S proteins that elicited polyfunctional CD4+ and CD8+ T cell responses (Fig. 4 and table S3). All mapped non-S proteins contained multiple immunogenic epitopes. A peptide was considered immunogenic, a “hit”, if in at least one patient, the percent of reactive cells was greater than the paired DMSO percent plus 3 times the standard deviation of all DMSO values (>DMSO+3SD) across the population. We investigated the novelty of these non-S immunogenic “hit” peptides by predicting the minimal epitopes for each “hit” binding to the responsive patients’ MHC alleles. A peptide was considered novel if none of its predicted binders were deposited in IEDB. Accordingly, we found that 42% of CD8+ and 23% of CD4+ T cell peptides we identified were novel, and the others were previously reported (fig. S8), providing additional confidence in the validity of our epitope discovery pipeline. Notably, certain immunogenic regions were commonly recognized by T cells across different convalescent patients (Fig. 4 and table S3). Together, these data demonstrate the wide breadth and robustness of T cell responses against non-S SARS-CoV-2 proteins and suggest that certain regions are immunodominant, significantly inducing T cell immunity across the population.

**Fig. 4.**
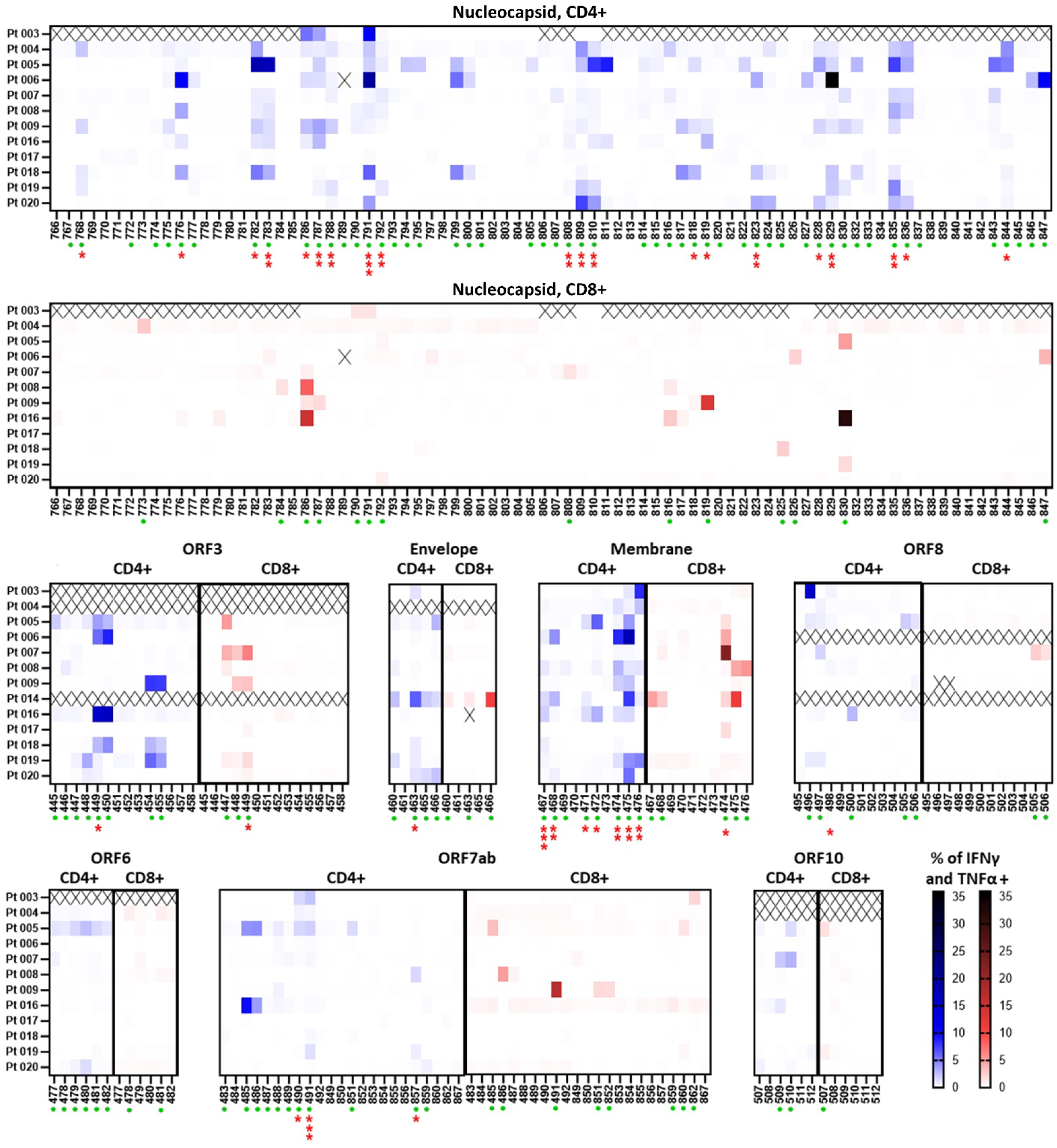
Deconvolution of T cell responses against non-spike SARS-CoV-2 proteins. T cells from convalescent, unvaccinated patients were expanded following stimulation with non-spike peptide pools. Then, expanded T cells were re-stimulated by individual 15mers constituting the peptide pools. Antigen-specific cytokine production by expanded T cell subsets, CD4+ (in blue) or CD8+ (in red), was measured by intracellular staining by flow cytometry. Heat maps demonstrate the percentage of reactive, polyfunctional T cells (secreting both IFN-γ and TNF-α) after normalization (subtraction of background, DMSO stimulation). y and x axes indicate the patients and peptides tested, respectively. Peptide sequences are reported in table S2. X indicates that the data was not collected. Statistical significance (p < 0.05) for peptides inducing T cell responses across the tested population was evaluated by Wilcoxon matched-pairs test comparing DMSO vs peptide stimulation and denoted in red stars if significant (immunodominant peptides). A peptide was considered immunogenic, a “hit”, if in at least one patient, the % of reactive cells was greater than the paired DMSO % plus 3 times the standard deviation (>DMSO + 3SD) of all DMSO values across the population, denoted by green dots.

### Conservation of non-spike immunodominant T cell epitopes

A high rate of recurrent mutations is observed across all regions of the SARS-CoV-2 genome (*45*), which can lead to escape from T cell immunosurveillance. Therefore, we examined the mutational diversity within non-S proteins, especially in immunodominant regions. First, we investigated the conservation of immunogenic (“hit”) sequences, listed in fig. 4, among SARS-CoV-2 variants by measuring the percentage of sequences that are deposited to GISAID and have an exact match to the “hit” peptides. We observed a high degree of conservation with a median over 99% for “hit” peptides inducing both CD4+ and CD8+ T cell responses (Fig. 5A). To more closely investigate the mutational diversity across the non-S peptidome, we computed the entropy of the observed amino acid frequencies on each codon position for each non-S protein utilizing sequencing and regional epidemiological count data obtained from GISAID EpiCoV database (*46*) and WHO Coronavirus dashboard (*47*), respectively, through different timepoints (*48*). We then compared the entropies of codons encoding experimentally validated immunogenic peptides (“hit” or immunodominant, table S3) vs those encoding the rest of the non-S proteome. We found that codons encoding immunogenic peptides, especially those that are dominant across the population, exhibit significantly lower entropy (Fig. 5B). To validate this, we conducted a parallel analysis utilizing entropy data from Nextstrain. Consistently, our findings confirmed the lower entropy values in immunogenic regions (fig. S9), thus further demonstrating that immunogenic non-S sequences are derived from mutationally constrained regions.

**Fig. 5.**
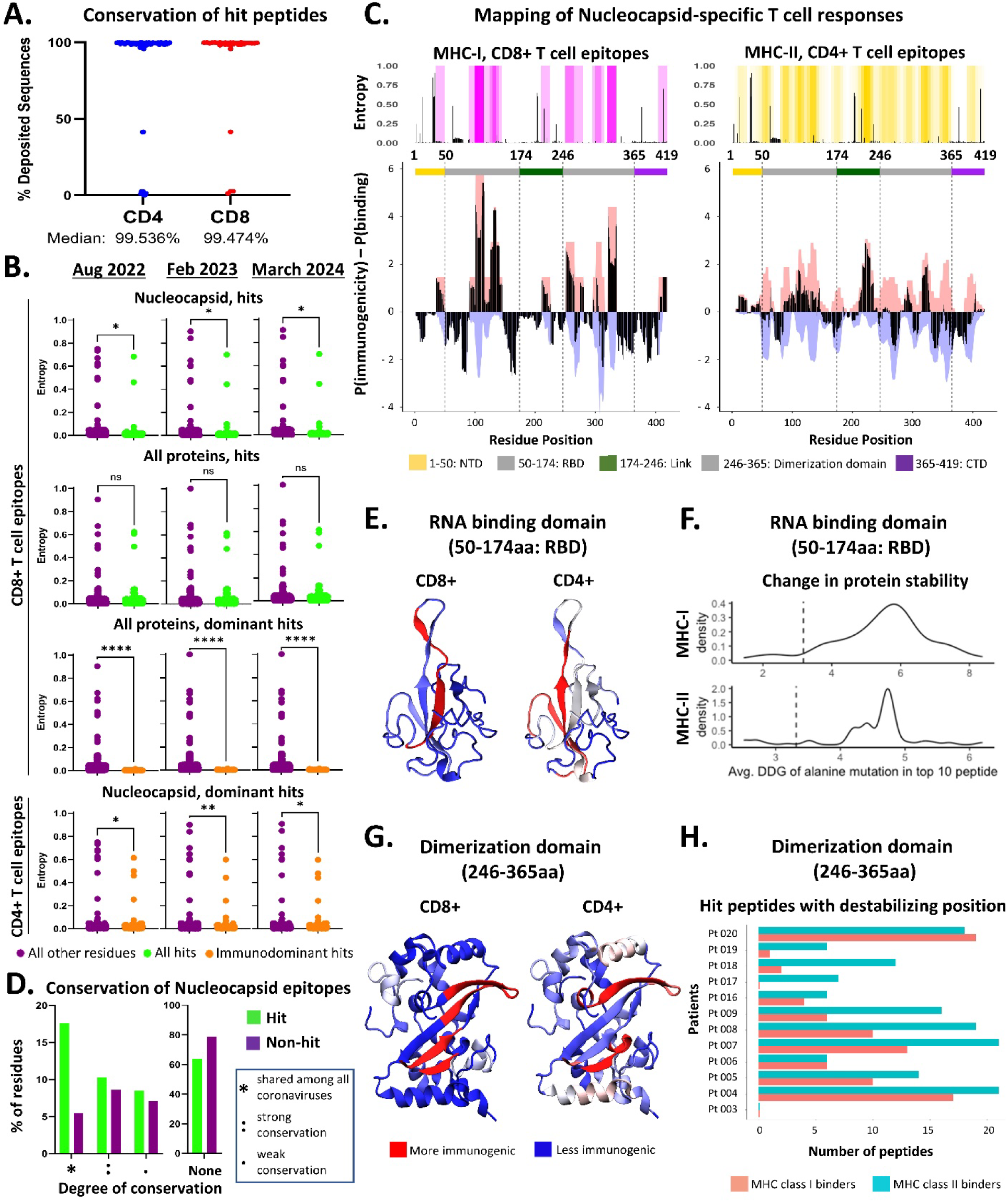
Conservation of T cell epitopes. **(A)** Conservation of the “hit” T cell peptides, identified in fig. 4, was measured as percent of previously deposited GISAID sequences with an exact match to the reactive peptide. The median conservation percentages for the CD4 and CD8 “hit peptides” were 99.54% and 99.47%, respectively. **(B)** Entropy values for the residues found in “hit” peptides (green) or “hit” peptides that were significantly enriched across the test population (orange) were compared to the entropy values for all other residues (purple) in the ORFs tested in our study. Statistical significance was calculated by Welch’s t-test. **(C)** Topography of immunogenic nucleocapsid residues is displayed. NTD: N-terminal domain, RBD: RNA binding domain, DD: dimerization domain, CTD: C-terminal domain. The black bars represent the difference between per residue immunogenicity (experimental, shaded in pink) and per residue binding value (predicted, shaded in lilac). Per residue immunogenicity values were generated by summing for each residue the normalized percentage of CD8 or CD4 T cells expressing both TNFα and IFNγ (>DMSO + 3SD) directed at each peptide for which the residue belongs. Per residue binding value corresponds to the frequency that a given residue appeared in a peptide predicted to bind to a patient MHC. The per residue values were scaled from 0-1 with 1 representing the residue with the highest immunogenicity value or the most frequently included in a predicted binding peptide and 0 representing the residue with the lowest immunogenicity or least frequent. The per residue values for binding predictions were then multiplied by −1 to reverse the sign. The Y axis was then transformed to represent the log odds ratio of the probability of being immunogenic vs predicted binders by dividing by the background probability (1/number of residues in the nucleocapsid protein). The normalized entropy per amino acid (aa) codon was also aligned with nucleocapsid, black bars denoting the degree of entropy and heatmaps showcasing the intensity of sharing of immunogenic residues in our cohort. **(D)** Summary of conservation degree for nucleocapsid CD8+ hit sequences analyzed by CLUSTAL O (1.2.4) multiple sequence alignment for H-COV Nucleocapsid Protein: 229E (UniProt Accession: A0A127AU35), NL63 (UniProt Accession: Q06XQ2), HKU1 (UniProt Accession: Q5MQC6), OC43 (UniProt Accession: P33469), SARS_2 (UniProt Accession: P0DTC9). Hit peptides are marked in green and other tested sequences are in purple. Sharing was denoted as the following: “*” Residue shared among all coronaviruses in sequence alignment, “:” Conservation between groups of strongly similar properties, “.” Conservation between groups of weakly similar properties. **(E)** Mapping of immunogenic regions within RNA binding domain (RBD) on the 6YVO crystal structure. **(F)** The average change in protein stability (DDG) of RBD upon mutating each residue to alanine for the top ten most immunogenic peptides. Positive values indicate destabilizing mutations. The dashed line indicates the sliding window average (length 9 for MHC-I and 15 for MHC-II) across the nucleocapsid RBD. **(G)** Mapping of immunogenic regions within the dimerization domain onto the 6ZWO crystal structure. **(H)** Number of immunogenic peptides that contained at least one residue that could destabilize the interface when mutated to the alanine position (DDG>0 when mutated) was calculated for each patient.

Next, we focused on nucleocapsid as it is the only non-S protein we epitope mapped across the entire protein sequence, and its immunogenicity data lacks any potential bias introduced by our initial peptide selection based on pMHC binding predictions. We aligned the normalized entropy values per amino acid residue with heatmaps displaying the extent of sharing of immunogenic residues across our cohort, alongside a map demonstrating regions enriched for immunogenic residues even despite the scarcity of predicted strong binders. This enrichment signified the production of immunogenic epitopes at a rate higher than would be suggested by the number of predicted binders in regions of lower entropy that also contained shared immunogenic residues (Fig. 5C). Importantly, these immunodominant, lower entropy regions mostly resided in highly structured domains of nucleocapsid protein, namely the RNA binding domain (RBD) and dimerization domain (DD) (Fig. 5C). This observation was not skewed by certain MHC alleles in our cohort preferentially binding to highly structured domains, as previously reported in other viral infections (*49*), since approximately 95% of all alleles were predicted to bind to epitopes from RBD and DD more frequently than other, less structured domains (fig. S10). Finally, the mapping of the immunogenic nucleocapsid epitopes reported in IEDB also showed an enrichment within RBD and DD compared to less structured regions (fig. S11).

Together these observations led to the hypothesis that immunogenic epitopes are enriched in the less diverse, functionally important regions of the SARS-CoV-2 proteome, thus likely exerting evolutionary constraints limiting mutational rates to avoid compromised viral fitness.

Accordingly, we examined epitope conservation throughout coronavirus evolution by comparing SARS-CoV-2 nucleocapsid amino acid sequence to other historical human coronaviruses, namely HCoV-229E, HCoV-NL63, HCoV-HKU1, and HCoV-OC43. Corroborating previous observations, immunogenic epitopes were more commonly found in fully conserved regions compared to non-immunogenic residues (Fig. 5D and fig. S12-13). To further investigate if the observed resistance to mutational pressure in the regions containing immunogenic epitopes is related to maintaining protein function, we conducted *in silico* analyses focusing on RBD and DD. First, we visualized the immunogenicity of each residue of RBD on its crystal structure and found that immunogenic regions mapped onto the internal beta sheet, particularly the β-hairpin (Fig. 5E), which is important for RNA binding (*50*, *51*). Then we tested whether the immunogenic residues were important for RBD stability by mutating each residue to alanine and comparing the energy of the modified structure to the wild type. The average change in protein stability calculated for the top ten most immunogenic peptides from each patient was greater than the overall average for residues across nucleocapsid RBD (Fig. 5F), indicating that most immunogenic peptides contain structurally important residues. Similarly, for DD, immunogenic regions mapped onto beta sheets (Fig. 5G), a critical motif in nucleocapsid oligomerization (*52*). Furthermore, alanine scanning revealed that mutations in immunogenic peptides from all patients, except Pt 003 which has missing data points, led to the destabilization of the protein-protein interface (Fig. 5H), impacting nucleocapsid function.

Taken together, the mutational diversity and biophysical analyses indicate that epitopes inducing T cell immunity reside in functionally and structurally important positions, potentially explaining the high conservation and lack of immune editing in immunogenic regions.

## DISCUSSION

Through the induction of adaptive immunity, vaccinations have markedly reduced SARS-CoV-2 infection rates, severe disease, and mortality. However, as antibodies wane over time and SARS-CoV-2 evolves to evade host immunity, especially neutralizing antibodies, it is necessary to develop new strategies that provide durable protection. Vaccine-induced S-specific T cell responses have been shown to be long-lasting (*17*), overall preserved against variants (*12*, *13*) and provide protection against COVID-19 even in the absence of antibody responses (*16*, *19*– *21*). Unlike antibodies, T cell targets are not limited to extracellular antigens, making non-S viral proteins a valuable source of potential T cell epitopes. Accumulating evidence demonstrates the immunogenicity of non-S proteins, as well as their role in mediating viral clearance and disease control (*22–25*, *38*, *53*, *54*). Thus, incorporating non-S proteins as targets in vaccine strategies could enhance durable protection against emerging variants. Accordingly, several preclinical studies have shown that inclusion of non-S antigens in immunization strategies provides robust protection against VOC (*55–58*). In a hamster model of SARS-CoV-2 VOC challenge, a dual S and N encoding mRNA vaccine, mRNA-S+N, had more robust control of Delta and Omicron variants in the lungs and upper respiratory tract than mRNA-S. Although less robust than the dual vaccine, mRNA-N alone was immunogenic and induced infection control (*57*). Moreover, a recently developed mRNA vaccine, BNT162b4, encoding segments of N, M and ORF1ab proteins, elicited T cell responses and provided protection against severe disease in murine models when administered alone or in combination with S encoding BNT162b2. Protection against VOC was enhanced with dual vaccination compared to vaccination with S encoding vaccine alone (*58*). BNT162b4 is currently being tested in a clinical trial in combination with the BNT162b2 Bivalent vaccine (NCT05541861).

As these new vaccines are designed, selection of epitope targets plays a critical role in achieving maximum efficacy and protection against VOC across a diverse human population. Strategies for selecting non-S targets may include utilizing entire proteins. Although this approach should unbiasedly generate a broad range of epitopes, the high number of targets may cause immune competition, potentially reducing the immunogenicity of key epitopes (*59*, *60*). Alternatively, non-S targets may be selected by utilizing computational algorithms predicting immunogenic epitopes. However, a high number of predicted epitopes does not elicit T cell responses (*61*). A thorough understanding of SARS-CoV-2-specific T cell responses, informing their breadth, immunodominance and conservation is needed to enable the design of broad-spectrum vaccines effective against emerging variants.

Here we studied SARS-CoV-2-specific T cell responses from vaccine recipients and convalescent patients via rigorous functional interrogation combined with *in silico* analyses. Our work revealed the (i) kinetics of vaccination-induced polyfunctional CD4+ and CD8+ T cell responses that remained intact against SARS-CoV-2 variants; (ii) immunodominant CD4+ and CD8+ T cell epitopes across SARS-CoV-2 proteome; and that (iii) the immunodominant epitope regions exist in protein domains of lower mutational diversity due at least in part to their functional importance. Together our findings identify critical non-S epitopes that are conserved and highly immunogenic across a diverse human cohort that can be utilized in future vaccine designs to elicit a focused T cell response effective against emerging VOC.

Our studies evaluating mRNA vaccine-induced adaptive immunity response kinetics showed that in SARS-CoV-2 infection-naïve individuals, while the antibody titers were peaked after the booster vaccine, prime vaccination was sufficient to induce T cell responses, which remained similarly high after the booster. This suggests that T cells may be key mediators of early response to vaccination. Accordingly, Oberhardt *et al.* showed that functional CD8+ T cells were mobilized even after 1 week of prime vaccination with BNT162b2 (*62*). Guerrera *et al.* also showed the induction of T stem cell memory after prime vaccination, correlating with durable T cell immunity (*17*). In our cohort the mRNA-1273 mRNA vaccine induced higher magnitude of T cell responses than the BNT162b2 vaccine. Similar differences between the two vaccines were reported for humoral responses (*34*). This is likely due different antigen loads and dosing schedules of the two vaccines. Overall, both vaccines induced polyfunctional T cells and similarly recognized tested VOC in *ex vivo* assays.

Our findings that while neutralizing antibodies diminish, T cell recognition of VOC is preserved in *ex vivo* assays agree with previous reports (*12*, *13*) and underline the critical role of T cell immunity in controlling emerging variants. This may be due to the wide breadth of T cell epitopes such that a reduced capacity of T cell recognition of certain mutated epitopes is compensated by the overall response. It may also be due to T cells that can cross-react with the mutated epitopes. Our findings support the relevance of both mechanisms. Our results with clonally enriched T cell populations specific to a single mutation region identified P26S and R246I as escape mutations, significantly evading recognition by CD4+ T cells. P26S was also previously reported to escape from CD4+ T cell responses (*13*). However, clonally enriched T cell populations specific to multiple mutation regions found in each VOC clade showed no difference in the overall recognition of mutant or ancestral sequences, compensating for any diminished response to a single epitope. In addition, modeling of T cell responses against ancestral and variant strains suggested cross-reactivity as a major predictive parameter of T cell immunity. This mathematical modeling approach we utilized only involved CD8+ T cell responses as it relied on peptide-MHC binding predictions, which perform poorly for MHC-II binding. However, it is also important to understand CD4+ T cell dynamics as SARS-CoV-2 primarily induces CD4+ T cell responses and the reduction in cross-recognition of S mutations in clonally enriched T cell populations was observed more robustly in the CD4+ T cell subset. Despite cross-reactivity being a key parameter of T cell response to VOC, including vaccination as an additional cross-reactivity measure with the complete Wuhan-1 S 9mers did not improve the performance of our model. This is likely because our dataset is not well suited to measure the vaccine’s effect since our experimental dataset is focused on a specific subset of anti-S T cell responses and *in vitro* expansion impacts clonal dynamics of T cells, reducing TCR diversity and allowing *in* vitro priming which may introduce a new set of TCRs.

Modeling of anti-S T cell immunity also identified peptide immunodominance as a key parameter. Others have also reported immunodominance, with a limited number of epitopes accounting for most of the total response against SARS-CoV-2 (*24*, *38*). Here we comprehensively mapped CD4+ and CD8+ T cell responses and found that patterns of immunodominance were similarly present in non-S proteins, a subset of epitopes robustly inducing T cell responses across a diverse cohort of patients. SARS-CoV-2 exhibits high rates of recurrent mutations across its genome (*45*), often as a mechanism to evade immune responses. This is observed particularly in VOC escaping from neutralizing antibody responses. Similarly, certain S mutations have been reported to cause escape from T cell recognition (*10*, *11*).

Consequently, immunodominant regions of the viral proteome may be expected to have high mutation rates to escape from T cell immunosurveillance. However, we found that non-S immunogenic epitopes were enriched in the less diverse regions of the SARS-CoV-2 proteome. Our data suggest that the high conservation and lack of immune editing in immunogenic regions is due to their positions in functionally and structurally important regions of the proteome, thus subject to evolutionary pressure restricting mutations and maintaining viral fitness. Another explanation is that MHC polymorphisms and diversity across populations provide an obstacle for viral immune evasion from T cell immunity.

Together, our findings provide a comprehensive dataset of functional T cell responses across MHC-typed study cohorts, which will help to greatly inform three ongoing research efforts in this field: i) technologies identifying SARS-CoV-2 epitopes and cognate TCRs to track antigen-specific T cell immunity (*63*, *64*) ii) T cell-based diagnostics and correlates of protection to complement antibody-based metrics and iii) design of next-generation coronavirus vaccines.

## MATERIALS AND METHODS

### Study Design

This study aimed to characterize SARS-CoV-2-specific T cell immunity elicited by vaccination or natural infection. We evaluated the kinetics of Spike-specific adaptive immunity induction following mRNA-based SARS-CoV-2 vaccination utilizing blood samples collected before vaccination, 14 days after 1st dose, 7 days after 2nd dose, and 14 days after 2nd dose. We also evaluated how Spike mutations present in variants of concern impact T cell recognition.

Furthermore, we investigated the breadth and magnitude of T cell responses against SARS-CoV-2 in unvaccinated, convalescent individuals. We mapped immunogenic T cell epitopes within non-spike proteins and evaluated their distributions across the viral proteome in the context of mutational diversity. SARS-CoV-2-specific antibody responses were measured in binding assays and pseudovirus neutralization assays. SARS-CoV-2-specific T cell responses were measured in *ex vivo* ELISPOT and antigen-specific T cell expansion assays utilizing overlapping peptides, 15mers with 5 amino acids offset. Sample sizes and statistical tests used were indicated in the figure legends and/or relevant method sections.

### Study Population

The study populations included individuals receiving initial two doses of mRNA-based COVID-19 vaccines, and two cohorts of unvaccinated, convalescent patients. The collection, processing and banking of vaccinated donor blood specimens were carried out by the Mount Sinai Human Immune Monitoring Core (HIMC). The sample collection for the first convalescent patient cohort (CPC; utilized in fig. 1E and 2B) was performed by the Cancer Immunotherapy Clinical Trials Team at Tisch Cancer Institute at Mount Sinai. PBMCs were isolated by density gradient centrifugation using Ficoll-Paque™ Plus (GE Healthcare) and cryopreserved in human serum containing 10% DMSO. The collection, processing and banking of blood samples from second convalescent patient cohort (ATLAS; utilized in fig. 3 and 4) were performed by the Mount Sinai Convalescent Plasma Donor Program. The use of patient-derived specimens was approved by the Institutional Review Boards at Mount Sinai and all patients provided written informed consent before the initiation of any study procedures. For ATLAS cohort a numerical COVID-19 disease severity scoring system was adapted from Moderbacher *et al.* (*14*) and indicated in the table S1 together with a summary of clinical demographics and sequence-based MHC-I/II genotyping (SBT) results (Histogenetics) for each subject. Although the infecting viral strains were not validated by sequencing, given the circulating SARS-CoV-2 strains during the time of infection, the convalescent patients in both cohorts are expected to have been infected with Wuhan-1.

Historical healthy donor specimens used as control were procured from New York Blood Center as leukopak prior to 2019 and PBMCs were isolated by density gradient centrifugation using Ficoll-Paque™ Plus (GE Healthcare). PBMCs were cryopreserved in human serum containing 10% DMSO.

### Peptide Selection and Synthesis

Custom libraries of overlapping peptides (OLPs) were chemically synthesized by GenScript and each peptide had at least 85% purity as determined by high-performance liquid chromatography. OLPs were typically 15mers and overlapped by 10 amino acids spanning the entire spike and nucleocapsid and selected regions from other SARS-CoV-2 proteins (Fig. 3A). To select regions in ORF1ab, ORF3a, envelope, membrane, ORF6, ORF7a, ORF7b, ORF8 and ORF10 proteins, we utilized a list of epitopes predicted to be immunogenic, published by Campbell *et al.* (*43*), and prioritized regions enriched in peptide-MHC (pMHC) interactions across different races (*44*) using IEDB’s population coverage tool (http://tools.iedb.org/population/help/#by_ethnicity).

This selection was supplemented with additional epitopes that had a predicted IC50 value less than 500 nM as determined using NetMHC 3.4. A complete list of all synthesized peptides and peptide pooling strategies are included in table S2.

### Pseudovirus Neutralization Assay

The neutralization capacity of patient plasma was assessed using a pseudotype particle (pp) infection system. Vesicular Stomatitis Virus (VSV) pseudotype particles were generated with a co-transfection strategy. This pseudovirus system was generated as described previously (*65*). A VSV[Rluc]-ΔG-G construct, which encodes the viral core genes with a renilla luciferase reporter substituted in place of the VSV entry surface glycoprotein gene, was used to infect 293T cells. These cells were also transfected with a plasmid encoding the full-length SARS-CoV-2 S from the codon-optimized Wuhan-Hu-1 isolate (NCBI accession no. NC_045512.2) and containing the D614G mutation. The resulting transfection product was a particle that encapsulated the viral RNA genome encoding the luciferase reporter and incorporated plasma-membrane expressed SARS-CoV-2 spike proteins. CoV2pp (VSVΔG-Rluc bearing the SARS-CoV-2 spike glycoprotein) at a concentration of ∼400 TCID50/mL was incubated with diluted, heat-inactivated patient plasma and then used to infect ACE2+ TMPRSS2+ 293T target cells. Following cell lysis (Promega), relative luminescence units (RLUs) were measured with a Cytation 3 reader. RLU values were normalized to those derived from cells infected with CoV2pp incubated in the absence of plasma. VSV-Gpp (VSVΔG-Rluc bearing the VSV-G entry glycoprotein) infection was used as a positive infection control and BALDpp (VSVΔG-Rluc bearing no protein) was used as a negative infection control. This assay has been validated in our laboratory using COVID-19 convalescent patient plasma and negative control plasma collected from healthy donors prior to the pandemic. The 4-point nonlinear regression curves were used to calculate ID50 values (GraphPad Prism v9) for each convalescent patient and vaccinated individual.

### Luminex Antibody Binding Assay

To detect antibody reactivity to the SARS-CoV-2 receptor binding domain (RBD) in individuals vaccinated with COVID-19 mRNA vaccines, we used a Luminex Binding Assay described previously (*66*). Briefly, SARS-CoV-2 mutant and wild-type (Wuhan-1 isolate) RBDs were covalently coupled to a uniquely labeled fluorochrome carboxylated xMAP bead set (Luminex) at 4.0 μg protein/million beads using a 2-step carbodiimide reaction with the xMAP Antibody Coupling Kit (Luminex). The coupled beads were pelleted, resuspended at 5×10^5^ beads/mL in storage buffer (PBS containing 0.1% bovine serum albumin (BSA), 0.02% Tween-20, and 0.05% sodium azide, pH 7.4), and stored at –80°C. The beads needed for a single run (2500 beads/well × number of wells) were diluted in assay buffer (PBS containing 0.1% BSA, 0.02% Tween-20) to a volume that delivered 2500 beads to each well in an aliquot of 50 μL/well. Serum/plasma was diluted in PBS, added as 50 μL/well to the wells containing the beads, and incubated at room temperature (RT) for 1 hour on a plate shaker at 600 rpm. After 2 washes with assay buffer, 100 μL/well of biotinylated antihuman total immunoglobulin (Abcam) at 2 μg/mL was added and incubated for 30 minutes at RT on a plate shaker. After 2 washes, 100 μL/well of streptavidin-PE (BioLegend) at 1 μg/mL was added and followed by a 30-minute incubation at RT on a plate shaker. After 2 additional washes, 100 μL of assay buffer/well was added and put on a shaker to resuspend the beads. The plate was read with a Luminex Flexmap 3D instrument.

### Enzyme-Linked Immunosorbent Assay (ELISA)

SARS-CoV-2 Spike-specific IgG antibody reactivity was assessed by ELISA. Briefly, 1 μg/mL of recombinant protein (RBD or Spike ectodomain) or 100 uM peptide was coated onto Nunc Maxisorp high-protein binding plates in PBS, overnight at 4°C. Plates were washed 6 times with washing solution (3 times with 1xPBS with 0.2% Tween20 and 3 times with 1X PBS) and incubated with blocking buffer (1xPBS with 5% dried milk powder) for 2 hours. After blocking, the plates were washed similarly, and patient serum was diluted with blocking buffer and left to incubate at room temperature (RT) for 2 hours. The plates were again washed prior to the addition of anti-human IgG conjugated to alkaline phosphatase (AP) at a 1:3500 dilution in blocking buffer. After 1 hour incubation at RT, the plates were developed with 0.6 mg/mL of substrate (AttoPhos, Promega) for 30 minutes and the developing reaction was stopped with 3M NaOH. The fluorescence was measured at 450nm (excitation)/555nm (emission) wavelength with an ELISA microplate reader (BioTek Synergy). The anti-SARS-COV-2 Spike monoclonal antibody CR3022 (Abcam) was used as a positive control for recombinant protein ELISAs.

### T Cell Immunogenicity Assays

SARS-CoV-2-specific T cell immunogenicity was evaluated by two methods: enzyme-linked immunosorbent spot (ELISpot) assays to measure *ex vivo* T cell responses and/or T cell expansion assays to robustly analyze peptide immunogenicity and elicited polyfunctional responses by CD8+ and CD4+ T cell subsets. For both assays, cryopreserved PBMCs were quickly thawed in 37°C water bath and transferred into RPMI medium (Thermo Fisher Scientific) containing DNase I (Sigma-Aldrich) at a final concentration of 2 U/mL, spun down and resuspended in media prior to being used in an assay described below.

For ELISpot assays, thawed PBMCs were resuspended in CTL-Test medium (ImmunoSpot) supplemented with GlutaMAX (Gibco). Cells were seeded at 2.5×10^5^ cells per well in at least duplicates in mixed cellular ester membrane plates (Millipore) which were previously (1-7 days) coated with 4 μg/mL of anti-IFN-γ antibody (clone 1-D1k, Mabtech) and blocked by incubating at least for 1 hour with RPMI medium containing 10% human serum. Cells were then stimulated with 1 μM of test peptides (custom peptide synthesis, GenScript) or control reagents as indicated in each relevant figure legend and costimulatory antibodies anti-CD28 (BD Biosciences) and anti-CD49d (BD Biosciences) added at a final concentration of 0.5 mg/mL. After 24 hours of incubation at 37°C, plates were processed for IFN-γ detection. Plates were first incubated with 0.2 mg/mL of biotinylated anti-IFN-γ antibody (clone 7-B6-1 by Mabtech) for 2 hours at 37°C, then 1 hour at room temperature (RT) with 0.75 U/mL streptavidin-AP conjugate (Roche) and lastly with the SigmaFast BCIP/NBT substrate for 15 minutes at RT. In between each step, plates were washed 6 times with PBS containing 0.05% Tween-20 and 3 times with purified water. Plates were scanned and analyzed using ImmunoSpot software.

For T cell expansion assays, a previously published protocol was utilized (*35*). Briefly, PBMCs were resuspended in X-VIVO 15 medium (Lonza) supplemented with cytokines promoting dendritic cell (DC) differentiation, GM-CSF (SANOFI, 1000 IU/mL), IL-4 (R&D Systems, 500 IU/mL) and Flt3L (R&D Systems, 50 ng/mL). Cells were seeded at 10^5^ cells per well in U-bottom 96-well plates and cultured for 24 hours before being stimulated with control reagents or pooled test peptides (custom peptide synthesis, GenScript), where each peptide was at a final concentration of 1 μM, together with adjuvants promoting DC maturation, LPS (Invivogen, 0.1 mg/mL), R848 (Invivogen, 10 mM) and IL-1β (R&D Systems 10 ng/mL), in X-VIVO 15 medium. Starting 24 hours after stimulation, cells were fed every 2-3 days with cytokines supporting T cell expansion, IL-2 (R&D Systems, 10 IU/mL), IL-7 (R&D Systems, 10 ng/mL) and IL-15 (Peprotech, 10 ng/mL) in complete RPMI media (GIBCO) containing 10% human serum (R10). After 10 days of culture, cells were harvested, pooled within groups, washed, resuspended in R10 and seeded in equal numbers into U-bottom 96-well plates. Expanded T cells were then re-stimulated with control reagents or 1 μM of test peptides, either pooled or individual, together with 0.5 mg/mL of costimulatory antibodies, anti-CD28 (BD Biosciences) and anti-CD49d (BD Biosciences), and protein transport inhibitors BD GolgiStopTM, containing monensin and BD GolgiPlugTM, containing brefeldin A. After 8 hours of incubation at 37°C, cells were processed for intracellular staining for flow cytometry using BD Cytofix/Cytoperm^TM^ reagents according to manufacturer’s protocol. The following antibodies were used: for surface staining CD3 (clone SK7, FITC), CD4 (clone RPA-T4, BV785) and CD8a (clone RPA-T8, APC) and for intracellular staining IFN-g (clone B27, PE), TNF-a (clone Mab11, PE/Cy7) and IL-2 (clone MQ1-17H12, BV605). All antibodies were purchased via BioLegend. LIVE/DEAD Fixable Blue Dead Cell Stain Kit by Thermo Fischer Scientific was used for live and dead cell discrimination. Data was acquired using the BD Fortessa and FlowJo V10 was used for analysis.

For both assays, DMSO (Sigma-Aldrich) was used at the equal volume of the test peptides and served as the vehicle/negative control. Unless noted otherwise all T cell immunogenicity data was shown after background normalization by subtracting the average of DMSO values (2 to 6 replicates per test group) from the corresponding spot numbers or cytokine+ cell frequencies following peptide stimulation. For peptide selection in Fig. 4 and 5, a peptide was considered immunogenic, a “hit”, if in at least one patient, tested the % of reactive cells was greater than the paired DMSO % average plus 3 times the standard deviation of all DMSO values across the population (>DMSO + 3SD).

### Modeling of T cell recognition and Model Selection

To model T cell responses (*m*) against spike mutations found in Alpha, Beta and Gamma variants (MT) or the corresponding WT (Wuhan-1) sequences in fig. 2C, we used a data likelihood approach. The complete set of experimental data measurements is represented as D={m_p_^Aκ,Bκ^:κ∈{Alpha,Beta,Gamma}; p∈patients; A,B∈{WT, MT}}, where A_κ_ stands for the pool (WT or MT) that was used for the first stimulation, and B_κ_ (also WT or MT) is the pool used for restimulation.

We proposed a mechanistic model to account for *m*_p_^*A_K_, B_K_*^ in terms of the putative immunogenicity of peptides included in the stimulation and restimulation pools. Due to the presence of multiple sources of noise, including experimental conditions and intrinsic differences in patient-specific immune responses, we assumed the experimental measurements to be drawn from an underlying Gaussian process with standard deviation σ and patient-specific immune amplitudes, *c*_*p*_. Our model prediction corresponding to *m*_p_^*A_K_,B_K_*^ is *c*_p_*R*(*B*_*K*_|*H*_p_, *A*_*K*_, Θ), where *R*(*B*_*K*_|*H*_p_, *A*_*K*_) is the T cell recognition of the restimulation pool, *H*_*p*_ is the patient’s set of 6 MHC class I alleles, and Θ the model parameters. The likelihood of observing the experimental data, *D*, can be expressed with the Gaussian (log) likelihood function,

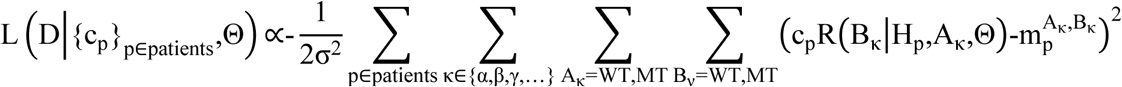

By maximizing the log-likelihood function we find the model parameters, Θ, which minimize the difference between our model predictions and the experimental data. The optimal patient-specific immune factors can be derived analytically by setting the partial derivative of the log likelihood function equal to zero and solving for *c*_p_, 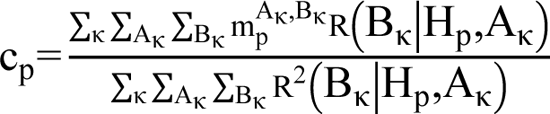. We used the Hyperopt package to optimize the remaining model parameters (*67*).

We considered T cell response as a two-step process: 1) peptide binding and presentation by MHC class I molecules (pMHC) and 2) binding of pMHCs by specific T cell receptors (TCRs). Accordingly, we computed T cell recognition at the 9mer level before aggregating over all 9mers in the peptide pools, and over the MHC-I set for each patient. The T cell recognition score for a single 9mer, *b*, from the restimulation pool *B*, in isolation from other 9mers in the pool, is computed as R(**b**,h)=p_MHC_(**a**,h,A)p_MHC_(**b**,h,B)exp[-βd(**a**,**b**)] The first term on the right-hand side accounts for the strength of presentation of *b* by an MHC molecule of allele 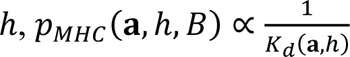. The second presentation term and the exponential term account for the cross-reactivity of peptides. The cross-reactivity distance, *d*(**a**, **b**), is a linear, sequence-based distance function derived from extensive epitope-TCR binding assays (*42*). In our equation we accounted for cross-reactivity of the corresponding WT-mutant peptides based on the assumption that 9mers present during the first stimulation (e.g., WT, *a*) which were well-presented would likely have elicited expansion of their cognate TCRs. Those expanded TCRs are then available to recognize restimulation pool 9mers (e.g.,Mutant, *b*), the binding strength depending on the extent of cross-reactivity with the first stimulation pool 9mers. The corresponding WT-mutant peptides only differed in most cases by a single amino acid; the cross-reactivity of the remaining pairs of peptides, depicted in fig. S3, are predicted to be negligible.

This basic model was extended with other components. Because all patients in this cohort were previously vaccinated, we can optionally include the vaccine as a zeroth stimulation event. This contributes another presentation term - for all spike protein 9mers, as taken from the sequence of the Wuhan-1 strain used in the original vaccine - and two additional cross-reactivity terms with the corresponding vaccine 9mer,

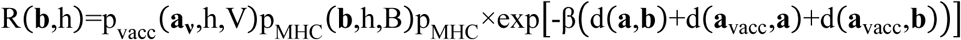

We can also include sequence similarity to immunogenic epitopes in the IEDB database (www.iedb.org). This approach has been previously used to define a TCR-immunogenicity score for neoantigenic peptides, attributing higher microbial peptide sequence similarity with increased TCR response (*40*, *41*). This introduces a new weight-parameter for the IEDB-similarity term, β_IEDB_,R(**b**|h)=p_pres_(**b**,h,B)*p_pres_(**a**,h,A)* exp[-β d(**a**, **b**)-β_IEDB_ d_min_(**b**,IEDB)], where d_min_(**b**,IEDB) is the distance of peptide **b** to the closest IEDB peptide.

We also tested different epitope dominance models to account for the contribution of different 9mers in the peptide pools. The immunodominance assumption is that the most immunogenic 9mer in a pool is largely responsible for the observed T cell response. Mathematically, this can be achieved by taking the maximum over all 9mer recognition terms. The maximum likelihood modeling framework was used to directly test the immunodominance hypothesis, and to compare it to a number of other assumptions using alternate aggregating functions including “sum”, where all 9mers contribute to the pool level response in proportion to their relative immunogenicity, “minimum”, not expected to be biologically realistic, and the “Boltzmann operator”, a parameterized smooth approximation to both max (α→∞) and min (α→-∞):

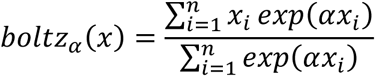

We assumed that all MHC-I alleles of a patient contribute to the pool-level response in a proportional manner, necessitating the summation as the aggregating function over *H*_*p*_, the set of 6 MHC-I alleles of the patient. The pool level response with 9mer-aggregating function *F* is therefore, R(B_κ_|H_p_,A_κ_,Θ)= ∑_h∈Hp_ F_b∈Bκ_ (R(**b**, h)) with model parameters Θ= {β,α}.

To compare the performance of these different models, we calculated the Bayesian Information Criterion (BIC), BIC=k log n −2 log L^^^, where *L*^^^ is the maximum log-likelihood score of a given model, *k* is the number of parameters of this model and *n* is the data size.

The recognition scores, R(**b**|ℎ), were evaluated for each subsequence 9mer **b** of the 15mer peptides in the pools, Alpha, Beta and Gamma with 161, 161, and 203 subsequences, respectively, for each MHC allele ℎ of the patients in the cohort. We computed the effective number of peptides that can be recognized among all the 9mers in a pool, the *perplexity*, *P = exp(H_rec_)*, with entropy the normalization constant assuring the entropy is computed over a probability distribution.

### Novelty of Immunogenic T Cell Epitopes

To determine the novelty of hit peptides, minimal epitopes binding to donor MHC alleles were predicted. For CD4 epitope analysis, 98 hit peptides (15mers, table S3) were included and predictions were performed using NetMHCIIpan 4.0. Epitopes (9-15mer) with a predicted binding affinity lower than 1000 nM were analyzed for their presence in IEDB. Similarly, for CD8 epitope analysis, 38 hit peptides (15mers, table S3) were included and predictions were performed using MHCflurry 2.0. Epitopes (9-11mer) with a predicted binding affinity lower than 500 nM were analyzed for their presence in IEDB. The determination of possible minimal epitopes was MHC-dependent, but the IEDB search was MHC-independent. The database was exported in August 2022. The MHC alleles of all responsive donors were included donors. A 15mer was considered novel if none of its predicted binders, across all patient MHC alleles, were present in IEDB. A permutation test was used to assess the significance of the novelty calculations for CD8+ T cell epitopes. Specifically, we generated synthetic lists of hits by random sampling without replacement from the full set of observations (peptides x patients). We did this 1000 times and ran the resulting lists of hits through the same pipeline to calculate the fractions that are novel. Using this resampling procedure, we found that the number of novel peptides we would expect to see by chance is significantly different (p<0.001) than what we observed.

### Conservation of T Cell Epitopes

To assess the conservation of selected immunogenic CD4+ and CD8+ T cell epitopes, the percentage of GISAID sequences deposited that have an exact match to the reactive peptide were measured. 100,000 random isolates from the GISAID database were subsampled and filtered to those with a unique accession number, where no proteins contain an unknown amino acid (X) after translation, have the full complement of all 27 proteins annotated, and were deposited during 2022. This analysis excluded ORF10 since it is not found in GISAID.

The degree of amino acid residue sharing throughout the human coronavirus family was determined by multiple sequence alignment using CLUSTAL O (1.2.4) for H-COV Nucleocapsid Proteins: 229E (UniProt Accession: A0A127AU35), NL63 (UniProt Accession: Q06XQ2), HKU1 (UniProt Accession: Q5MQC6), OC43 (UniProt Accession: P33469), SARS_2 (UniProt Accession: P0DTC9).

Peptide binding predictions in fig. 5C and S10 were performed using netMHCpan-4.1 and netMHCIIpan-4.1 for MHC-I and MHC-II predictions, respectively. Peptide binding thresholds were set to 2% for MHC-I and 10% for MHC-II (*68*).

### Diversity Analysis

To assess mutational diversity during global circulation in the SARS-CoV-2 genome, entropy of the observed amino acid frequencies was computed. The sequence and regional epidemiological count data were combined to account for regional disproportions in sequencing efforts while computing amino acid frequencies. The sequence data used was obtained from the GISAID EpiCoV database (*46*) available until 2024-03-21. For quality control, the 3’ and 5’ regions of sequences were truncated and sequences that contain more than 5% ambiguous sites or have an incomplete collection date were removed. All sequences were aligned against a reference isolate from GenBank 80 (MN908947), using MAFFT v7.525 (*69*). Weekly infection rates were used for individual countries as reported from the WHO Coronavirus dashboard (*47*) (download date: 2024-03-21). Infection counts were distributed equally over all days within a given reporting week. Individual countries were grouped into coarse grained geographical regions *r* by their continents (North America, Europe, Asia, South America, Africa and Oceania). The sequence counts in each geographical region *r* and day *t* were computed as *N*_*r*_(*b*) = ∑_*i* ∈*r*_ *t* (*t* − *t*_*i*_), where *ω* (τ) = exp(−τ^4^/4σ^2^) with σ = 11 days is the smoothing kernel function. Combining the sequence counts with the corresponding incidence data, *I*_*r*_(*t*), weight factors *m*_*r*_(*t*) = *I*_*r*_(*t*)/*N*_*r*_(*t*) measuring the incidence per sequence count in each region were recorded (*9*, *48*). Each GISAID sequence collected at region *r* and on date *b* was weighted by a factor *m*_*r*_(*b*)/ ∑_*r, t*_ *m*_*r*_(*r*) when computing the amino acid frequencies *x*(*a*, *i*) on individual codon positions *i*. The entropy for position *i* was then computed as *H*(*i*) = − ∑_*a*_ *x*(*a*, *i*) *log x* (*a*, *i*).

To determine where experimentally validated immunogenic T cell epitopes reside, the entropy value for each codon (and its corresponding residue) plotted as a bar graph was superimposed with an immunogenicity value for each individual residue across the nucleocapsid protein. Per residue immunogenicity values were calculated as the sum of the reactive T cell percentages against each peptide spanning that residue.

As an alternative approach, Shannon entropy data from Nextstrain available in August 2022 and February 2023 (https://nextstrain.org/ncov/gisaid/global) were analyzed for each codon. These entropy values were calculated using complete viral genome sequences deposited to the Global Initiative on Sharing All Influenza Data (GISAID) from around the world. This approach was not applied to analysis by April 2024 since the most recent GISAID datasets did not include codon/residue entropies.

### Structural Analysis of Nucleocapsid

To assess whether immunogenic T cell epitopes contained structurally important residues, immunogenic regions were mapped onto 6YVO and 6ZWO crystal structures, RNA binding and dimerization domains of nucleocapsid, respectively. The change in protein stability (DDG) was measured by mutating each residue of the protein into alanine and comparing the energy of that structure to the wildtype. This resulted in a DDG value being assigned to every residue with positive values indicating destabilizing mutations and negative values indicating stabilizing mutations. The top ten most immunogenic peptides were isolated from each patient, and the average DDG for those peptides was determined by averaging the values of individual residues. Alanine scanning was performed using the Rosetta analysis suite (*70*). The impacts on monomer stability of the RBD were probed using the cartesian_ddg application (*71*).The interface of the dimerization domain was analyzed using the Robetta alanine scanning server of the 6ZWO crystal structure (*72*).

### Statistical Analysis

Statistical tests used are indicated in the figure legends and in relevant methods sections. Briefly, statistical analyses were performed using GraphPad Prism versions 8 and 9. All error bars represent mean + SD unless noted otherwise in the figure legend. To evaluate significance, Wilcoxon matched-pairs test was used for paired samples and Welch’s t-test was used for unpaired samples. Significance was denoted by *, **, ***, ****, indicating P < 0.05, P < 0.01, P < 0.001, and P < 0.0001, respectively.

## Supplementary Materials

Figs. S1 to S13

Tables S1 to S3

## Acknowledgments

We gratefully acknowledge all data contributors, i.e., the Authors and their Originating laboratories responsible for obtaining the specimens, and their Submitting laboratories for generating the genetic sequence and metadata and sharing via the GISAID Initiative, on which a portion of this research is based.

We thank Vincenza Itri and Susan Zolla-Pazner at the Department of Medicine, the Division of Hematology and Medical Oncology, Icahn School of Medicine at Mount Sinai, New York, NY, USA, for generously allowing use of their Luminex machine in their laboratory for multiplexed antibody binding assays and providing technical supervision of reagent and equipment use.

We thank the members of the Mount Sinai COVID-19 Biobank Team listed here for their contributions to patient sample collection and processing: Charuta Agashe, Priyal Agrawal, Alara Akyatan, Kasey Alesso-Carra, Eziwoma Alibo, Kelvin Alvarez, Angelo Amabile, Carmen Argmann, Kimberly Argueta, Steven Ascolillo, Rasheed Bailey, Craig Batchelor, Noam D. Beckmann, Aviva G. Beckmann, Priya Begani, Jessica Le Berichel, Dusan Bogunovic, Swaroop Bose, Cansu Cimen Bozkus, Paloma Bravo, Mark Buckup, Larissa Burka, Sharlene Calorossi, Lena Cambron, Guillermo Carbonell, Gina Carrara, Mario A. Cedillo, Christie Chang, Serena Chang, Alexander W. Charney, Steven T. Chen, Esther Cheng, Jonathan Chien, Mashkura Chowdhury, Jonathan Chung, Phillip H. Comella, Dana Cosgrove, Francesca Cossarini, Liam Cotter, Arpit Dave, Travis Dawson, Bheesham Dayal, Diane Marie Del Valle, Maxime Dhainaut, Rebecca Dornfeld, Katie Dul, Melody Eaton, Nissan Eber, Cordelia Elaiho, Ethan Ellis, Frank Fabris, Jeremiah Faith, Dominique Falci, Susie Feng, Brian Fennessy, Marie Fernandes, Nataly Fishman, Nancy J. Francoeur, Sandeep Gangadharan, Daniel Geanon, Bruce D. Gelb, Benjamin S. Glicksberg, Sacha Gnjatic, Joanna Grabowska, Gavin Gyimesi, Maha Hamdani, Diana Handler, Jocelyn Harris, Matthew Hartnett, Sandra Hatem, Manon Herbinet, Elva Herrera, Arielle Hochman, Gabriel E. Hoffman, Jaime Hook, Laila Horta, Etienne Humblin, Suraj Jaladanki, Hajra Jamal, Jessica S. Johnson, Gurpawan Kang, Neha Karekar, Subha Karim, Geoffrey Kelly, Jong Kim, Seunghee Kim-Schulze, Edgar Kozlova, Arvind Kumar, Jose Lacunza, Alona Lansky, Dannielle Lebovitch, Brian Lee, Grace Lee, Gyu Ho Lee, Jacky Lee, John Leech, Lauren Lepow, Michael B. Leventhal, Lora E. Liharska, Katherine Lindblad, Alexandra Livanos, Bojan Losic, Rosalie Machado, Kent Madrid, Zafar Mahmood, Kelcey Mar, Thomas U. Marron, Glenn Martin, Robert Marvin, Shrisha Maskey, Paul Matthews, Katherine Meckel, Saurabh Mehandru, Miriam Merad, Cynthia Mercedes, Elyze Merzier, Dara Meyer, Gurkan Mollaoglu, Sarah Morris, Konstantinos Mouskas, Emily Moya, Naa-akomaah Yeboah, Girish Nadkarni, Kai Nie, Marjorie Nisenholtz, George Ofori-Amanfo, Kenan Onel, Merouane Ounadjela, Manishkumar Patel, Vishwendra Patel, Cassandra Pruitt, Adeeb Rahman, Shivani Rathi, Jamie Redes, Ivan Reyes-Torres, Alcina Rodrigues, Alfonso Rodriguez, Vladimir Roudko, Panagiotis Roussos, Evelyn Ruiz, Pearl Scalzo, Eric E. Schadt, Ieisha Scott, Robert Sebra, Hardik Shah, Mark Shervey, Pedro Silva, Nicole W. Simons, Melissa Smith, Alessandra Soares Schanoski, Juan Soto, Shwetha Hara Sridhar, Stacey-Ann Brown, Hiyab Stefanos, Meghan Straw, Robert Sweeney, Alexandra Tabachnikova, Collin Teague, Ryan Thompson, Manying Tin, Kevin Tuballes, Scott R. Tyler, Bhaskar Upadhyaya, Akhil Vaid, Verena Van Der Heide, Natalie Vaninov, Konstantinos Vlachos, Daniel Wacker, Laura Walker, Hadley Walsh, Wenhui Wang, Bo Wang, C. Matthias Wilk, Lillian Wilkins, Karen M. Wilson, Jessica Wilson, Hui Xie, Li Xue, Nancy Yi, Ying-chih Wang, Mahlet Yishak, Sabina Young, Alex Yu, Nina Zaks, Renyuan Zha.

## Author contributions

Conceptualization: CCB, MB, VR, BDG, ML, NB Formal Analysis: CCB, MB, MT, EW, TO Funding Acquisition: BDG, NB

Investigation: CCB, MB, LV, MT, EW, TO, DG, YB, DR, PK, JK, VR, DH Methodology: CCB, MB, LV, MT, EW, TO, NV, ML, NB

Resources: KYO, KDS, GK, HA, NK, CM, RG, KN, DDV, DD, DR, JS, PSG, AC, MM, SKS, BL, AW, VS

Supervision: CCB, DC, NV, ML, NB Visualization: CCB, MB, MT, EW, TO Writing – Original Draft: CCB, MB, MT, EW

Writing – Review & Editing: CCB, MB, MT, EW, TO, YB, DH, KDS, GK, HA, NK, RG, DD, EC, PSG, BL, VS, DC, ML, NB

## Data and materials availability

The findings of a portion of this study (Fig. 5A-D) are based on 8,897,424 individual genome sequences and associated metadata published in GISAID’s EpiCoV database (*46*) up to the following dates 2022-08-21, 2023-02-26, and 2024-03-21 via EPI_SET_240918oy. All sequences in this dataset are compared relative to hCoV-19/Wuhan/WIV04/2019 (WIV04), the official reference sequence employed by GISAID (EPI_ISL_402124). To view the contributors of each individual sequence with details such as accession number, Virus name, Collection date, Originating Lab and Submitting Lab and the list of Authors, visit 10.55876/gis8.240918oy. All other data needed to evaluate the conclusions of the paper are available in the main text or the supplementary materials. Python scripts used for model fitting are available upon request.

